# Reelin deficiency contributes to long-term behavioral abnormalities induced by chronic adolescent exposure to Δ9-tetrahydrocannabinol in mice

**DOI:** 10.1101/2020.08.10.245449

**Authors:** Attilio Iemolo, Aisha Nur, Patricia Montilla-Perez, Victoria B Risbrough, Francesca Telese

## Abstract

Heavy and frequent use of cannabis during adolescence increases the risk of developing psychiatric disorders. However, the neurobiological mechanisms underlying this vulnerability remain largely unknown. Here, we explore whether adolescent vulnerability to long-term behavioral effects of cannabis is modulated by *Reelin*, a gene implicated in the development of the brain and of psychiatric disorders. To this aim, heterozygous Reeler (HR) mice, that express reduced level of *Reelin*, were chronically exposed during adolescence to high doses (10mg/kg) of Δ9-tetrahydrocannabinol (THC), a major psychoactive component of cannabis. Mice were tested in early adulthood with multiple behavioral assays, including working memory, social interaction, locomotor activity, anxiety-like responses, stress reactivity, and pre-pulse inhibition. Compared to wild-type (WT), HR mice treated with THC showed impaired social behaviors, elevated disinhibitory phenotypes and increased responsiveness to aversive situations, in a sex-specific manner. Independent of THC exposure, HR mice also spent more time exploring unfamiliar objects, indicating that Reelin modulates novelty seeking behavior. To identify the neuronal ensemble underlying this elevated novelty seeking in HR mice, we mapped the regional brain expression of the immediate early gene, *Fos*, in mice exposed to novel objects. HR mice exhibited reduced neuronal activation in the lateral septum, a subcortical brain structure implicated in emotions, cognition and reward processes. Overall, these findings show that (1) *Reelin* deficiency influences behavioral abnormalities caused by heavy consumption of THC during adolescence, and (2) that Reelin plays a role in the neurobiological mechanisms underlying disinhibitory behaviors, such as novelty seeking.

**Significant Statement:** The link between cannabis abuse and the development psychiatric disorders, especially in adolescents, makes understanding the neurobiological mechanisms underlying cannabis effects on the brain a significant biomedical problem. Reelin is a key signaling molecule in the development of the adolescent brain and of psychiatric disorders, but its role in modulating the behavioral changes induced by cannabis remain unknown. Here, we report an interaction between *Reelin* deficiency and chronic adolescent exposure to THC, a major psychoactive component of cannabis. This interaction led to cognitive deficits, disinhibitory behaviors and altered emotional reactivity in mice, in a sex-specific manner. These experiments are the first to establish a link between Reelin signaling and the endocannabinoid system targeted by THC.

## Introduction

Heavy and frequent cannabis use by adolescents has been linked epidemiologically to increased risk of developing psychiatric conditions, including schizophrenia, psychosis, and substance use disorders (Volkow, 2016). Similarly, animal studies show that administration of synthetic cannabinoids (e.g. THC) during adolescence perturbs a wide range of behaviors, including memory, social interaction, anxiety, and sensorimotor gating by targeting the endocannabinoid (eCB) system (Rubino et al., 2015). Despite the evidence of possible detrimental health outcomes associated with adolescent cannabis use, a recent survey in the US revealed a substantial increase in daily use of cannabis and a decreased perception of the risks associated with its regular use by adolescents (Johnston, 2018). Further, the increasing legalization of recreational cannabis use has led to calls to understand whether such policies put adolescents at higher risk of developing psychiatric disorders. All this emphasizes the need to better understand the neurobiological mechanisms associated with heavy consumption on cannabis during adolescence (Wilkinson et al., 2016).

During adolescence, the brain undergoes continuous remodeling of its structure, connectivity, and plasticity (Sturman and Moghaddam, 2011; Arain et al., 2013). In addition, substantial hormonal changes during adolescence influence not only reproductive functions, but also the emergence of sex differences in cognitive, social, and emotional behaviors (Schulz and Sisk, 2016). Thus, the adolescence is considered a critical period wherein brain development may be altered by the exposure to psychoactive drugs, which can lead to sex-specific behavioral abnormalities and increased risk for psychopathology in adulthood (Cousijn et al., 2018; Lisdahl et al., 2018).

However, the neurobiological mechanisms underlying the increased vulnerability of the adolescent brain to the effects of THC remain poorly understood. The goal of this study is to examine the potential role of Reelin signaling in modulating the behavioral effects of cannabis on the adolescent brain.

Reelin is a protein of the extracellular matrix that is predominately expressed in neuronal cells and plays a key role in brain development and synaptic plasticity (D’Arcangelo et al., 1995). During embryonic stages, Reelin activates an extensive signaling cascade that is critical for the proper migration and cell positioning of cortical neurons (Sekine et al., 2014). During adolescence, Reelin signaling promotes the development of the synaptic excitation/inhibition (E/I) balance within the prefrontal cortex (Iafrati et al., 2014; Bouamrane et al., 2016). In the mature brain, Reelin is required for learning and memory by regulating the N-methyl-D-aspartate receptors (NMDA-R) function and the expression of neuronal activity-dependent genes (Weeber et al., 2002; Qiu et al., 2006; Niu et al., 2008; Rogers et al., 2011; Telese et al., 2015). In humans, *Reelin* deficiency has been linked to the development of psychiatric disorders (Ishii et al., 2016). Thus, the HR mice, that expressed lower level of Reelin, have been proposed as a valid animal model to study neurodevelopmental psychiatric disorders (Lossi et al., 2019). Whether there is a functional relationship between *Reelin* deficiency and the consequences of early-life exposure to high levels of THC remains unknown.

To this aim, we examined the long-lasting behavioral outcomes of chronic adolescent exposure to high doses of THC (10mg/kg) in female and male HR mice. We compared HR mice to their WT littermate controls in a battery of behavioral tests exploring different facets of cognitive and emotional responsiveness, including working memory (Sannino et al., 2012), social interaction (Yang et al., 2011), anxiety-like responses (Bailey and Crawley, 2009), stress reactivity (Can et al., 2012), and pre-pulse inhibition (Geyer et al., 2002). To further characterize the neuronal ensembles underlying behavioral differences between WT and HR mice, we examined the expression of the immediate early gene, *Fos*, in four brain regions that were activated after exposure to unfamiliar objects. This is the first study to investigate the relationship between *Reelin* deficiency and the effect of adolescent exposure to THC in mice.

## Materials and Methods

### Animals

All experimental procedures were approved by the institutional animal care and use committee at University of California, San Diego. Mice were housed (3-4 per cage) under a 12h light/12h dark cycle and provided with food and water ad libitum. HR mice were bred in house using the B6C3Fe a/a-Relnrl/J line (The Jackson Laboratory, #000235), which carry a null mutation in the Reelin gene (D’Arcangelo et al., 1995). HR express ~40% reduced level of Reelin mRNA in brain tissues (**Suppl. Fig. 1A-B**).

### Drug treatment protocol and experimental design for behavioral analysis

THC was provided by the U.S. National Institute on Drug Abuse and was dissolved in a vehicle solution consisting of ethanol, tween, and 0.9% saline (1:1:18) on the day of administration. Vehicle or THC (10 mg/kg) were administered daily to adolescent mice by intraperitoneal injections from post-natal day (PND) 28 to PND 48, which cover the adolescent period in mice (Laviola et al., 2003) (**Suppl. Fig. 1C**). This dose was based on previous studies using low to high ranges of THC in mice (Trexler et al., 2018; Kasten et al., 2019). Moreover, this dose used has been shown to elevate plasma levels of THC in rodents (> 100 ng/mL) similar to those found in humans smoking cannabis (Huestis, 2007; Zuurman et al., 2008; Nguyen et al., 2016). To examine the long-term effects of chronic adolescent exposure to THC, mice were tested 15 days after a drug abstinence period, starting at PND 63. We used 9 cohorts of mice to perform multiple behavioral assays. Cohorts 1 to 5 were subjected to locomotor activity, open field (OF), six-different objects (6-DOT), light-dark (LD), three-chamber social approach, and tail suspension (TS) tests. Cohorts 6 to 9 were subjected to locomotor activity, acoustic startle response (ASR) and pre-pulse inhibition (PPI) tests. Mice were tested between 10:00 am and 5:00 pm. Behavioral assays were conducted on separate days and all behavioral tests were performed once on each mouse. The number of mice used in each behavioral assay is reported in Table 1. To prevent bias due to olfactory cues, the behavioral apparatus was cleaned with diluted ethanol solution in between mice.

**Table 1.**
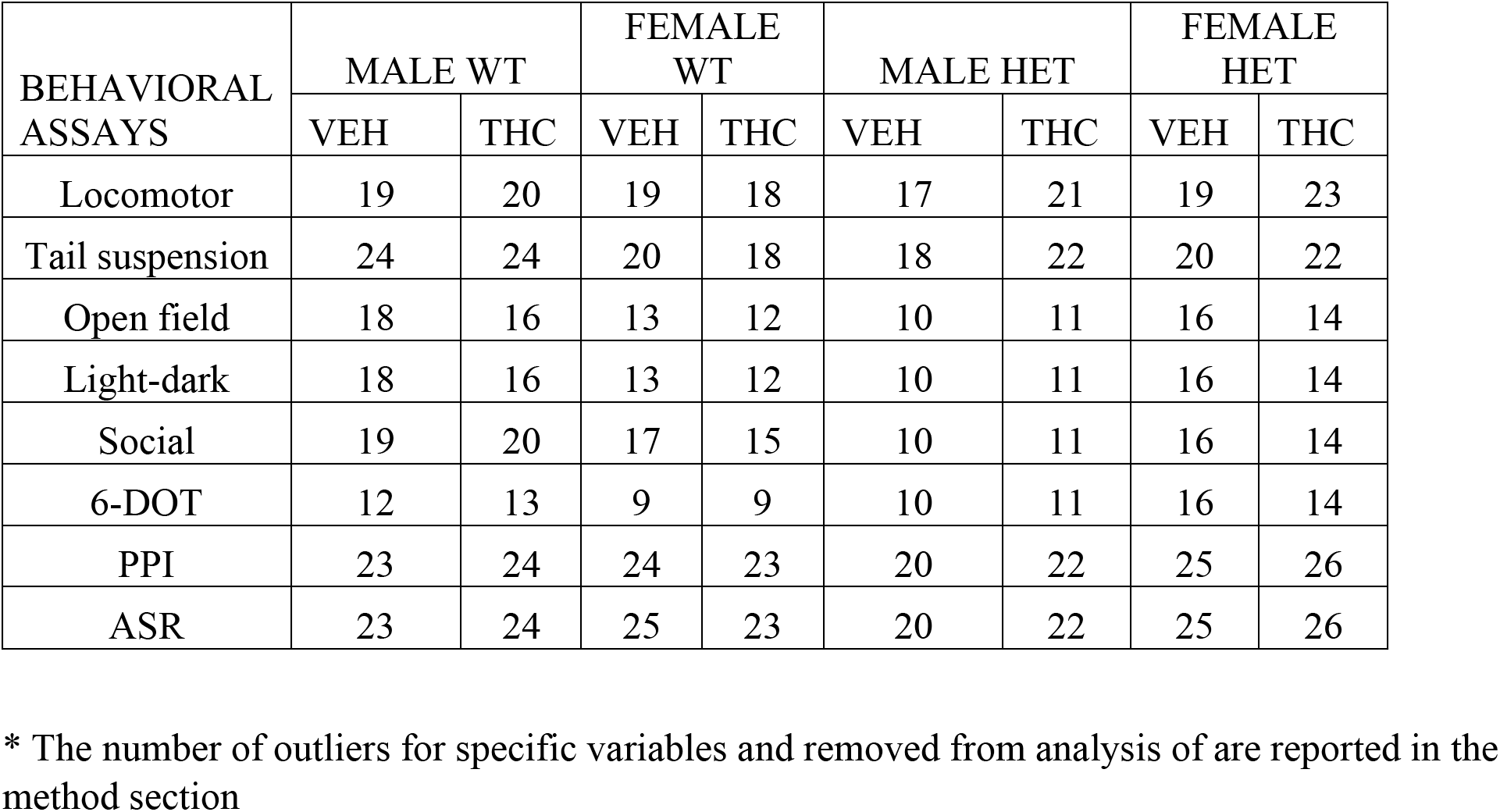
Number of mice used for behavioral analysis*

| BEHAVIORAL<br>ASSAYS | MALE WT |  | FEMALE<br>WT |  | MALE HET |  | FEMALE<br>HET |  |
| --- | --- | --- | --- | --- | --- | --- | --- | --- |
|  | VEH | THC | VEH | THC | VEH | THC | VEH | THC |
| Locomotor | 19 | 20 | 19 | 18 | 17 | 21 | 19 | 23 |
| Tail suspension | 24 | 24 | 20 | 18 | 18 | 22 | 20 | 22 |
| Open field | 18 | 16 | 13 | 12 | 10 | 11 | 16 | 14 |
| Light-dark | 18 | 16 | 13 | 12 | 10 | 11 | 16 | 14 |
| Social | 19 | 20 | 17 | 15 | 10 | 11 | 16 | 14 |
| 6-DOT | 12 | 13 | 9 | 9 | 10 | 11 | 16 | 14 |
| PPI | 23 | 24 | 24 | 23 | 20 | 22 | 25 | 26 |
| ASR | 23 | 24 | 25 | 23 | 20 | 22 | 25 | 26 |

\* The number of outliers for specific variables and removed from analysis of are reported in the method section

### Six different objects test

We used the 6-DOT to study short-term working memory in mice, as previously described (Sannino et al., 2012). First, mice were left free to explore an empty arena (60×40×35 cm) during a 10 min habituation trial. Afterwards, mice were exposed to six different objects for a total time of 10 min (familiarization trial). After an inter-trial interval of 1 minute, mice were exposed to identical copies of the familiar objects, but one object was substituted with a novel object (test trial). The exploration time across different trials was measured with Anymaze and was used to calculate a discrimination index (DI) as follows: (time spent exploring novel object – average time spent exploring familiar objects) / Total time spent exploring novel + familiar objects. The time exploring the novel object and the discrimination index (DI) were used as indexes of novelty-induced exploratory activity and working memory, respectively.

### Three-chamber social approach test

The three-chamber social approach test was used to measure social behaviors, as previously described (Yang et al., 2011). The apparatus comprised of three-chambered box with dividing walls with small openings to allow free exploration of the three chambers, such as one empty central chamber, one side chamber containing an empty small wire cage (novel object) and one side chamber containing a stranger mouse inside a small wire cage (novel mouse). The target mouse was first placed in the center chamber and allowed to explore the apparatus for 15 min. After introducing the novel mouse and the novel object in the side chambers, the mouse was allowed to explore for 10 min. The placement of the novel mouse or novel object in the left or right chambers was systematically alternated in between trials. The time spent in each compartment and the time spent actively sniffing the novel mouse or the novel object were manually scored. Longer time spent with or exploring the novel mouse versus the novel object was considered an index of sociability. Sociability index (SI) was calculated as [(time spent exploring or sniffing novel mouse - time spent exploring or sniffing novel object) / (total time spent exploring or sniffing novel mouse and novel object)].

### Tail suspension test

The TS test was used to measure motor responses under aversive conditions (Can et al., 2012). Mice were suspended by their tails with tape to a bar in a position that they could not escape or hold on to nearby surfaces. The test lasted for 6 minutes and the mobility time (sec) was manually scored for each minute. We analyzed was the mobility time (sec) as the sum of the final 5 minutes.

### Open field test

The OF test was used to measure anxiety-like behavior(Bailey and Crawley, 2009). Mice were placed randomly in one of the 4 corners of an open plexiglass arena (60×40×35 cm) for 5 minutes. The total time spent in and the latency to entry the center of the arena (s) were recorded and scored using Anymaze.

### Light-dark test

The LD test was used to measure anxiety-like behavior(Bailey and Crawley, 2009). The animals were tested for 10 min in a light–dark rectangular box (60×40×35 cm) in which the aversive light compartment (40×40×35 cm) was illuminated by a 100 lux light. The dark side (20×40×35 cm) had an opaque cover and ~ 0 lux of light. The two compartments were connected by an open doorway, which allowed the subjects to move freely between the two compartments. The test began by placing the animal in the dark compartment. The time spent in the light compartment (s), the latency to enter in the light chamber (s), and the total number of transitions, were measured using Anymaze.

### Acoustic startle response and pre-pulse inhibition test

The ASR and PPI tests were used to assess stress reactivity and sensorimotor gating functions. The tests were performed with a startle reflex measuring apparatus (SR-LAB; San Diego Instruments, San Diego, CA), as previously described (Toth et al., 2013). The system comprises a piezoelectric unit that transduces vibrations into signals when mice startle inside the plexigas cylinder. First, mice are placed in a plexiglas cylinder with background noise (65 decibel [db]) for 5 min (acclimation phase). Then, mice were subjected to a total of 179 trials, for a total of 25 min, including: (a) startle trials (40 milliseconds [ms] with 80, 90, 100, 110 and 120 db acoustic pulses), (b) prepulse+startle trials: 20 ms with acoustic prepulses of 3 (68), 6 (71) and 12 (77) db above background noise followed, 100ms later, by a 40 ms 120-db startling pulse. Startle amplitude was measured every 1ms over a 65ms period beginning at the onset of the startle stimulus. Average startle amplitude over the sampling period was taken as the dependent variable. Percent PPI at each pre-pulse intensity was calculated as 100 - [(startle response for prepulse/startle response for startle-alone trials) x 100].

### Locomotor activity

Locomotor activity was measured using the video tracking system Anymaze (Ugo Basile, Varese, Italy). Mice were placed in an empty open field (60×40×35 cm) for 20 minutes. The distance traveled (m) was recorded in 5 minutes intervals and used as an index of locomotor activity in a novel environment.

### Body weight measurements

Body weight (g) was measured through the course of the drug administration protocol and at PND63 when the behavioral assays began. The change in body weight was calculated as the difference between body weight at any given day and body weight at PND 28.

### RNA Extraction, cDNA Synthesis and qPCR

Brain tissue from PFC *n* = 8 female WT, *n* = 6 male WT, *n* = 6 female HR, *n* = 5 male HR mice was homogenized in TRIzol Reagent (#15596018, Thermo Fisher Scientific) and Zirconium beads (#Zr0B05-RNA, Next Advance) using the Bullet Blender homogenizer (BBX24B, Next Advance,). RNA was extracted on columns with the Direct-Zol RNA miniprep kit (#R2051, Zymo Research). To quantify *Reelin* expression levels, equal amounts of cDNA were synthesized using the SuperScript VILO MasterMix (#11755-050, Thermo Fisher Scientific) and mixed with the qPCRBIO SyGreen Blue Mix (#17-507DB, PCR Biosystems) and 5 pmol of both forward (5’-GGACTAAGAATGCTTATTCC -3’) and reverse (5’-GGAAGTAGAATTCATCCATCAG -3’) *Reelin* primers. ACTB was amplified as an internal control (5’-ATGGAGGGGAATACAGCCC -3’) and reverse (5’-TTCTTTGCAGCTCCTTCGTT -3’).

### Immunohistochemistry (IHC)

To examine Fos expression by IHC in WT and HR mice, we used *n* = 6 male WT, *n* = 6 male HR, *n* = 4 female WT, and *n* = 9 female HR littermates. Mice were subjected to the 3 trials of the 6-DOT test, as described above. To confirm that the exposure to the behavioral task triggers Fos expression, a separate group of 4 WT mice was used, including *n* = 1 mouse for each trial and *n* = 1 mouse without any behavioral exposure (home cage). Thirty minutes after the last trial, mice were anesthetized with isoflurane and fixed via transcardial perfusion with 4% paraformaldehyde (PFA) in phosphate buffered saline (PBS). Brain tissues were processed as previously described (Iemolo et al., 2020). Briefly, 30-μm thick coronal slices were stained with a primary antibody recognizing Fos (1:1000, ABE457, EMD Millipore, RRID:AB_2631318) and a secondary anti-rabbit antibody (donkey anti-rabbit Alexa Fluor 488 (1:1000, #A21206, Thermo Fisher Scientific). Images were acquired with a fluorescent microscope (BZX800, Keyence Corporation, Osaka, Japan). Different brain areas were identified using a mouse brain atlas as a reference (Paxinos G, 2001). For the medial prefrontal cortex (mPFC), 5 sections from bregma + 2.46 to + 1.10 mm, ± 340 μm apart, were stained and 4 images for each section were captured for each mouse. For the CA1 and dentate gyrus (DG) subregions of the hippocampus (HP), 4 sections from bregma −1.46 to - 2.46 mm, ± 250 μm apart, were stained and 4 images for each section were captured for each animal. For the lateral septum (LS), 1 section from bregma 1.42 mm to 1.10 mm was stained and 1 image for each section was captured for each mouse. The number of Fos+ cells was counted by two examiners blind to the experimental conditions. All numbers are presented as mean ± SEM.

### RNA fluorescent in situ hybridization (FISH)

To examine the expression of various transcripts in specific cell types of the lateral septum, we performed RNA FISH using the RNAscope Multiplex Fluorescent Reagent Kit v2 (ACD, #323100) and following the instructions for ‘fixed-frozen tissue sample’ of the user manual (ACD, USM-323100). Tissue sections corresponding to the LS (bregma 0.26-0.50 mm) from *n* = 1 mouse exposed to the 3 trials of the 6-DOT, as described above, were hybridized with a mix of three probes; Reelin (ACD, # 405981) + Slc32a1 (ACD, #319191-C3) + Chat (ACD, # 408731) or Fos (ACD, #316921) + Slc32a1 + Chat. LSc and LSr were identified using the Allen Mouse Brain Atlas. To assess both tissue RNA integrity and assay procedure, a separate group of sections were incubated with probes (data not shown). Images were acquired with the Keyence fluorescent microscope.

### Statistical analysis

To examine how factors (treatment, genotype, sex) affected mice behavior or Fos expression, we used linear mixed models (LMM) in JMP pro v. 15.0 (SAS Institute, Inc). LMM allow modeling of both fixed and random effects (which subsume repeated measures) (Quinn GP, 2002; Zuur AF, 2016). We incorporated the categorical predictor variables of interest as fixed effects and included all possible interactions. We included mouse cohorts and individual subjects as random effects to account for possible non-independence of the data. We ensured assumptions of approximate normality and variance homogeneity were met by inspecting plots of residuals versus predicted values, and by inspecting quantile-quantile plots with 95% confidence limit curves. When residual plots indicated that it was appropriate for repeated measures, we used a covariance structure that allowed variances to differ across the levels of the repeated variable (Garrett M. Fitzmaurice, 2011). We analyzed the untransformed data in all but few traits where a fourth root transformation mitigated variance heterogeneity (e.g. latency to light, latency to center, Vmax of acoustic startle, body weight change). When significant interactions were found, planned post-hoc pairwise comparisons were performed to identify differences among specific genotype and treatment groups. We report *P*-values that remained significant after controlling for multiple comparisons by holding the ‘false-discovery rate’ to 0.05 using the Benjamini-Hochberg method (Hochberg, 1995). Outliers were detected using the Huber M-estimation method (Huber, 1973) in JMP pro v. 15.0 and removed when appropriate (*n* = 3 in latency to light, *n* = 2 latency to center, *n* = 2 time spent sniffing, *n* = 1 sociability index, *n* = 1 in % PPI).

### Factor analysis of behavioral assays

Factor analysis is a data reduction method for understanding underlying relationships among variables (Bartholomew, 2008). We performed factor analysis using the variables from multiple behavioral assays, such as 6-DOT (DI, total exploration time in test trial [T3]), social interaction (time sniffing novel mouse, difference time sniffing novel mouse vs novel object), OF (time in center), LD (time in light), TS (mobility time). Factor analysis was computed in JMP 15.0 Pro and was conducted with varimax rotation with a factor-loading cutoff of 0.3 (Manly and Navarro Alberto; Stephens, 1996). The number of factors retained in our model was selected by inspecting the ‘elbow’ on the scree plot curve with factors retained if their eigenvalues were greater than 1 (Cattell, 1966). With these settings, a three-factor model was generated for our dataset. The loadings of the observed variables on the extracted factors are shown in Fig. 6. Factor scores for individual mice were extracted and used as variables for subsequent LMM analysis to identify their associations with treatment, sex, and genotype.

## Results

### Working memory was impaired after chronic adolescent exposure to THC, and in male HR mice

To assess the long-term effects of chronic adolescent exposure to THC on working memory in WT and HR mice, we used a modified novel object recognition test that uses 6 instead of 2 objects (6-DOT) (Sannino et al., 2012; Olivito et al., 2016). This test evaluates recognition memory under conditions of high loads of information processing, which is referred to as memory span and is considered a form of working memory. A discrimination index (DI) was calculated to determine the amount of time mice explored the novel object compared to the familiar objects. A preference for the novel object is considered as a sign that mice remember the familiar objects (Sannino et al., 2012). Higher rates of exploratory activity in response to the novel object is also interpreted as a sign of novelty seeking behavior (Flagel et al., 2014).

Chronic THC exposure during adolescence decreased the DI (working memory) of mice across all groups by ~31% compared to the vehicle-treated control group (**Fig. 1A-B**, treatment effect, F_1,86_ = 16.4, *P* < 0.0001). HR mice also showed decreased DI, but the magnitude of the genotype effect varied among sexes (**Fig. 1C**, sex x genotype interaction, F_1,86_ = 5.7, *P* = 0.019). Compared to WT, female HR mice did not show impaired working memory, but male HR mice showed a decreased DI that was similar to the level of impairment caused by THC treatment in male WT mice (**Fig. 1C**, paired *t* test, *t* = 2.9, *P_adj_* = 0.019).

**Figure 1:**
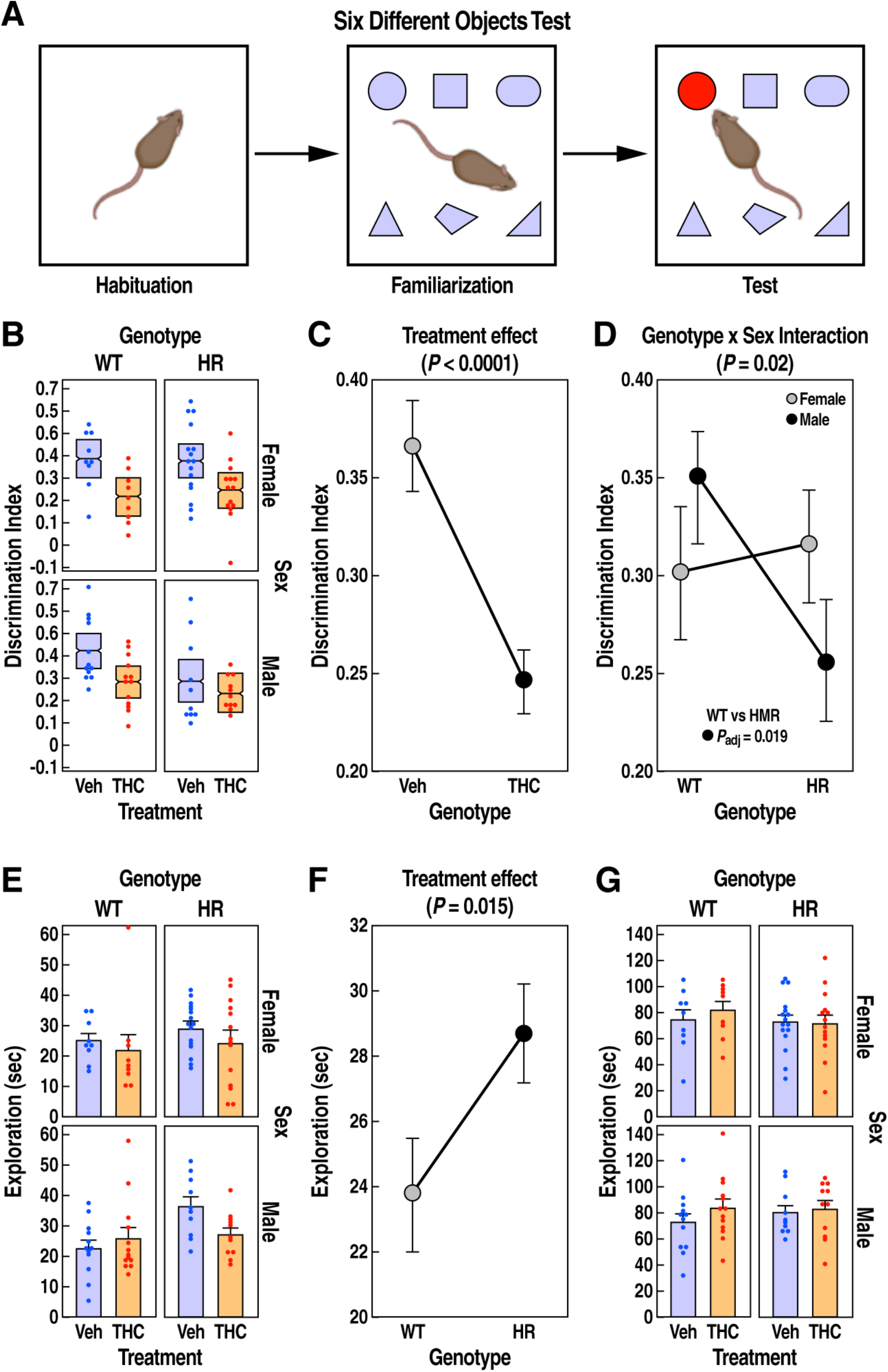
Six-different objetcs test in WT and HR after chronic adolescent exposure to THC. **(A)** 6-DOT with habituation, familiarization, and test trials. **(B)** Discrimination index (DI) is shown as mean ± 95% confidence intervals. **(C)** THC reduced the DI (mean ± SEM) across all groups (main treatment effect, *P* < 0.0001, LMM). **(D)** Male HR mice showed a reduced DI (mean ± SEM) compared to male WT mice (sex x genotype interaction, *P* = 0.02, LMM). **(E)** Exploration time (s) in test trial is expressed as mean ± SEM. **(F)** HR mice spent longer time (mean ± SEM) exploring the objects in the test trial (main genotype effect, P = 0.016, LMM). **(G)** Exploration time (mean ± SEM) in familiarization trial did not change across groups. Adjusted *P* < 0.05 for planned post-hoc pairwise t tests are reported.

This assay also revealed that HR mice displayed a ~25% increase in novelty-induced exploratory activity compared to WT controls (**Fig. 1D-E**, genotype effect, F_1,84_ = 6.1, *P* = 0.015). This effect was triggered by the novel object, as it was revealed during the test phase while the total exploratory activity toward objects did not vary among groups in the initial familiarization phase (**Fig. 1F**, main effects and interactions *P*s > 0.3).

To further evaluate potential confounding effects of general locomotion or body weight having an impact on exploratory activity in this task, we analyzed changes in spontaneous locomotor activity in an empty arena or in body weight in WT and HR mice (**Suppl. Fig. 2A-B**). There was no effect of THC treatment, genotype, or sex on locomotor activity or body weight at the time of testing, suggesting that the deficits in working memory observed in WT and HR mice were not driven by changes in locomotion or body weight.

### Social behaviors were impaired by chronic adolescent exposure to THC in HR male mice

To assess the long-term effects of chronic adolescent exposure to THC on social interaction behavior in WT and HR mice, we used the three-chamber social approach test in which a mouse is given the choice to spend time interacting with a novel mouse or a novel inanimate object, which are referred to as “exploration targets” in our analysis (Yang et al., 2011). We considered two aspects of social interaction: (1) time spent in the chamber with the novel mouse versus the novel object, and (2) time actively sniffing the novel mouse versus the novel object, which is a more specific measure of social interaction (Yang et al., 2011). Higher time spent in the chamber with or sniffing the novel mouse compared to the novel object was considered an index of “sociability”.

All mice explored the novel mouse more than the inanimate object, confirming that the behavioral assay can detect sociability across all groups (**Fig. 2A-2D**, exploration target effect, *F*_1,119_ = 49.5, *P* < 0.0001; *F*_1,113_ = 83.3, *P* < 0.0001, respectively). There were no significant effects of treatment, genotype and sex on time spent in the chamber (*P* > 0.1). In contrast, when analyzing time sniffing, the effect of chronic adolescent exposure to THC varied among the two exploration targets (**Fig. 2E**, treatment x exploration target interaction, F_1,113_ = 4.1, *P* = 0.046). Precisely, THC had no significant effect on time spent sniffing the novel inanimate object compared to vehicle (**Fig. 2E**, paired *t* test, *t* = 0.06, *P_adj_* = 1); however, THC caused a ~20% decrease in time spent sniffing the novel mouse compared to vehicle across all groups (**Fig. 2E**, *t* = 3.1, *P_adj_* = 0.003). The suppressive effect of THC on social interaction depended on the genotype (**Fig. 2F**, treatment x genotype interaction, *F*_1,108_ = 6.1, *P* = 0.015); a post-hoc analysis revealing that only HR mice treated with THC exhibited a significant reduction in overall time sniffing when compared to vehicle treated HR mice (paired *t* test, *t* = 3.2, *P_adj_* = 0.006). However, when a sociability index was calculated, we did not detect any effect of treatment or genotype (all effects and interaction *Ps* >0.1, **Suppl. Fig. 3A-B**).

**Figure 2:**
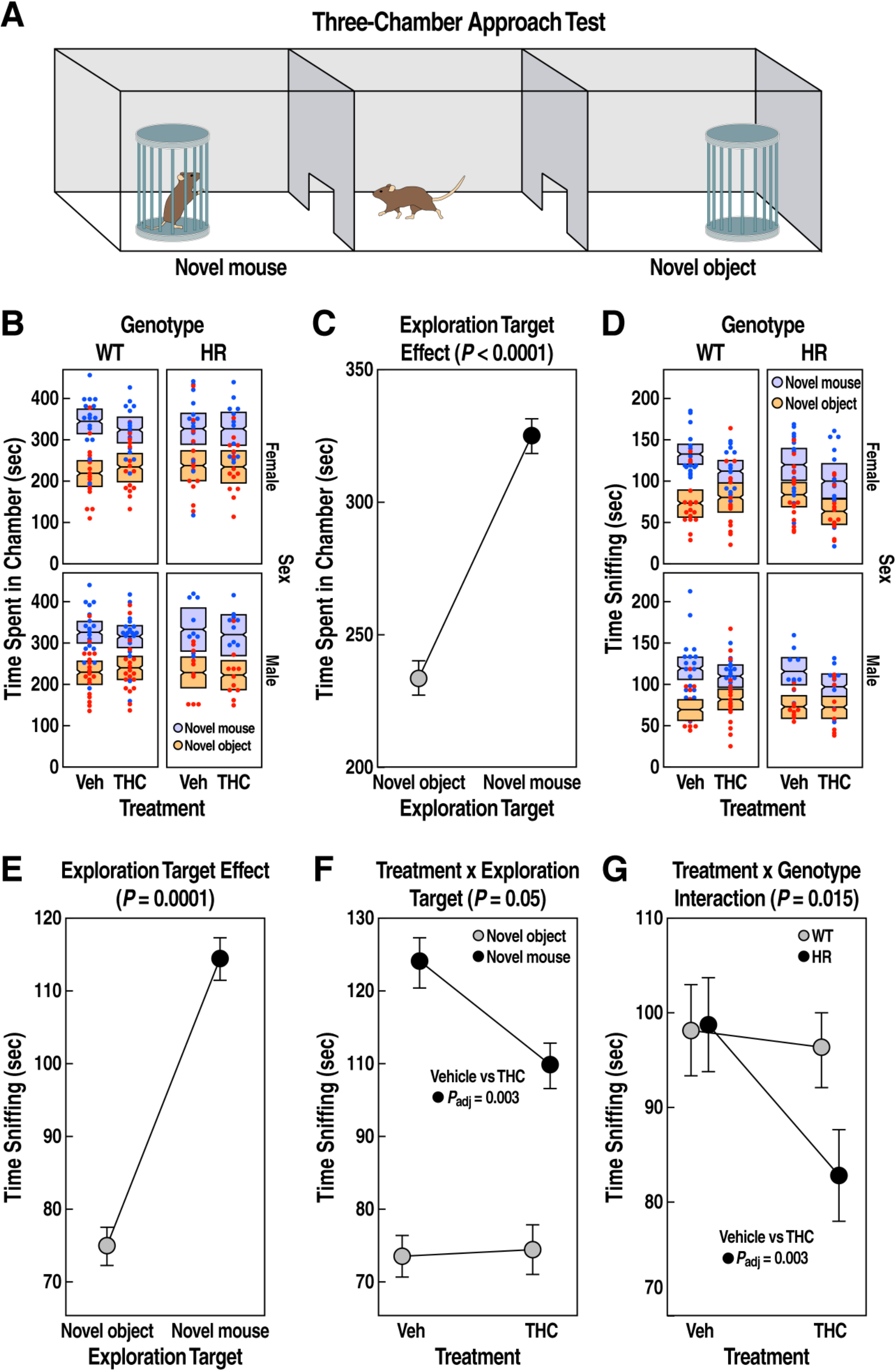
Social interaction behavior in WT and HR after chronic adolescent exposure to THC. **(A)** Three-chamber interaction test diagram. Time (s) spent in chamber **(B)** or sniffing **(D)** is expressed as mean ± 95% confidence intervals. **(C)** Mice spent more time (mean ± SEM) in the chamber or **(D)** more time sniffing (mean ± SEM) the novel mouse compared to the novel object (main exploration target effect, *P* < 0.0001, LMM). **(E)** THC reduced social investigation (post-hoc pairwise *t* test *P*_adj_ = 0.003; treatment x exploration target interaction, *P* = 0.05, LMM). **(G)** HR exposed to THC during adolescence showed reduced social interaction behavior (post-hoc pairwise *t* test *P*_adj_ = 0.006; treatment x genotype interaction, *P* = 0.015, LMM).

### Chronic adolescent exposure to THC increased anxiety-like behavior in female HR mice

To evaluate how impaired Reelin signaling influences anxiety-like behavior in mice chronically exposed to THC during adolescence, we compared WT and HR mice in the LD and OF tests. These behavioral assays examine anxiety-like responses by which animals avoid illuminated or open areas (Crawley and Goodwin, 1980; Bailey and Crawley, 2009). Decreased anxiety (increased disinhibition) is represented by mice spending more time in the light or open compartments, and by having a shorter latency to enter the brightly lit or center chambers.

In the LD test, THC increased the overall time spent in the light compartment (**Fig. 3A**, treatment effect, *F*_1,102_ = 8.5, *P* =0.0043), but the strength of these effects varied among genotypes and sexes (**Fig. 3B**, treatment x genotype x sex interaction, *F*_1,97.5_ = 4.4, *P* = 0.038). This effect was mainly driven by female HR mice treated with THC, which spent 43% longer time in the light compared to the vehicle-treated group (**Fig. 3B**, paired *t* test, t = 3.6, *P_adj_* = 0.004). A separate analysis revealed that the latency to enter the light chamber was also influenced by genotype and sex (**Fig. 3D-E**, sex x genotype interaction, F_1,47_ = 5, *P* = 0.0197). Female HR mice showed ~18% increase in latency compared to female WT, indicating an increased baseline anxiety-like responses (paired *t* test, *t* = 2.5, *P_adj_* = 0.03). However, the latency to enter the light compartment was not significantly influenced by THC (treatment effect, *F*_1,97_ = 0.02, *P* = 0.88).

**Figure 3:**
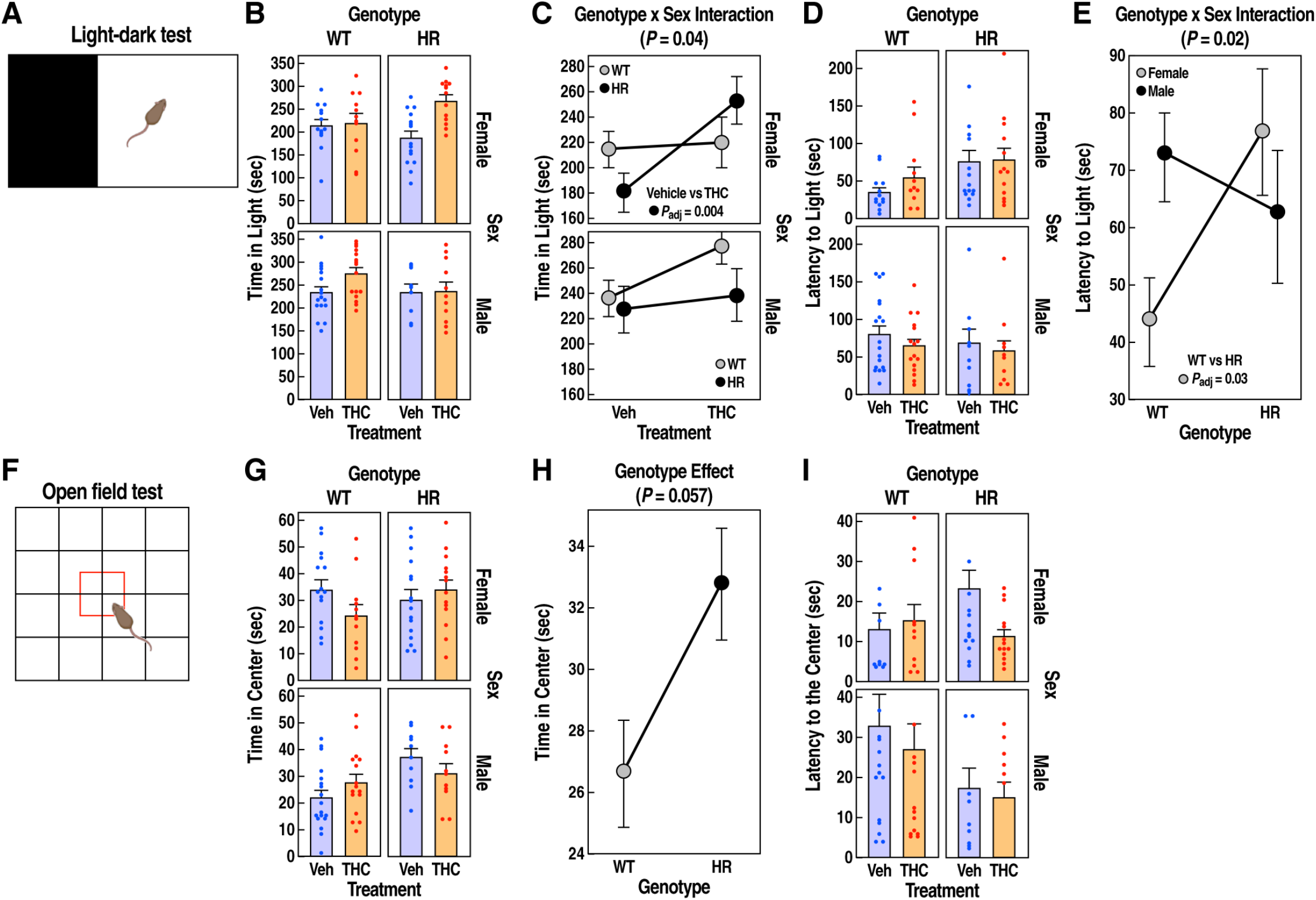
Anxiety-like responses in WT and HR following chronic adolescent exposure to THC. **(A)** Light-dark test schematic. **(B)** Time (s) spent in the light compartment is expressed as mean ± SEM. **(C)** Female HR treated with THC during adolescence showed reduced anxiety (treatment x genotype x sex interaction, *P* = 0.04, LMM). **(E)** Female HR mice showed increased latency (s) to enter the light compared to WT controls (genotype x sex interaction, *P* = 0.02, LMM). **(F)** Open field test schematic. **(G)** Time (s) spent in the center of the arena is expressed as mean ± SEM. **(H)** HR mice showed reduced anxiety (main genotype effect, *P* = 0.057, LMM). **(I)** Latency (s) to enter the center compartment is expressed as mean ± SEM and did not change across groups. Adjusted *P* < 0.05 for planned post-hoc pairwise t tests are reported.

In the open field test, HR mice spent ~18% more time in the center (**Fig. 3F-G**, genotype effect, *F*_1,102_ = 3.7, *P* = 0.057 marginally significant), suggesting a reduced anxiety-like behavior. Time spent in center also varied among sexes and treatment groups (treatment x genotype x sex interaction, *F*_1,99_ = 6.1, *P* = 0.016); however, despite this significant overall variability, a post-hoc analysis did not identify significant differences between specific pairs of groups. Latency to enter the open area did not vary among groups (**Fig. 3H**, all *P* > 0.2).

### Male HR mice showed enhanced ability to strive against stress following chronic adolescent exposure to THC

To test the effect of adolescent exposure to THC on reactivity to aversive conditions in WT and HR mice, we performed the TS test, an assay commonly used to screen anti-depressive drugs and to measure behavioral despair in mice (Can et al., 2012). An increased mobility has been associated with enhanced ability to strive against stress and is broadly related to active coping, impulsive and aggressive behaviors (Strekalova et al., 2004; Brockhurst et al., 2015).

We quantified total mobility time in a 6-minute tail suspension assay in WT and HR mice. THC treatment influenced total mobility time, but the effect varied among sexes and genotypes (**Fig. 4A**, treatment x sex x genotype interaction, *F*_1,158_ = 12.8, *P* = 0.0005). While THC did not significantly change the mobility time of either male or female WT mice compared to the vehicle controls (paired *t* test, *t* = 1.9, *P_adj_* = 0.07; *t* = 0.5, *P_adj_* = 0.6, respectively), it induced a 19% increase in mobility time for male HR mice (**Fig. 4B**, paired *t* test, *t* = 2.6, *P_adj_* = 0.04). We also observed sex differences in baseline mobility time for which female HR mice showed a 21% increase in mobility time compared to male HR mice (**Fig. 4B**, paired *t* test, *t* = 3, *P_adj_* = 0.03).

**Figure 4:**
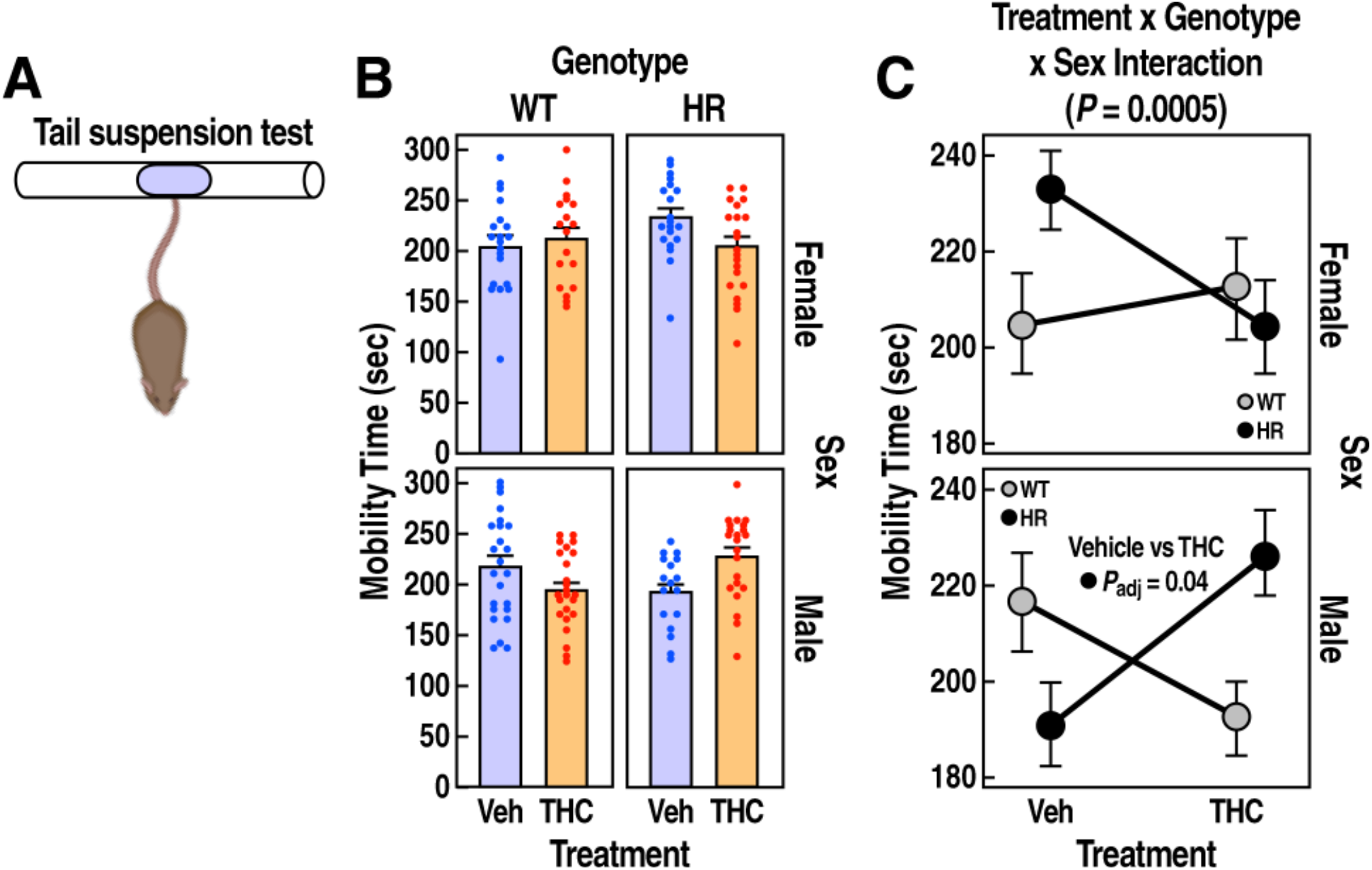
Reactivity to aversive conditions in WT and HR mice following chronic adolescent exposure to THC. **(A)** Tail suspension test diagram. **(B)** Mobility time (s) is expressed as mean ± SEM. **(C)** Male HR mice treated with THC during adolescence showed increased mobility (treatment x genotype x sex interaction, *P* = 0.0005, LMM). Adjusted *P* < 0.05 for planned post-hoc pairwise t tests are reported.

### Reelin deficiency led to enhanced startle responses and pre-pulse inhibition

To evaluate psychotic-like behaviors in WT and HR mice following chronic adolescent exposure to THC, we quantified the ASR and the % PPI. The ASR is a reflexive reaction to a sudden acoustic stimulus (Swerdlow et al., 2001; Geyer et al., 2002). The % PPI is used as a measure of sensorimotor gating, which occurs when a weak pre-pulse stimulus suppresses the response to the subsequent startling stimulus. PPI is impaired in schizophrenia patients and serves as an animal model of schizophrenia (Cadenhead et al., 1993).

We first examined the differences in ASR of WT and HR mice at various acoustic intensities (65, 80, 90, 100, 110, 120 db, **Fig. 5A**). THC did not influence ASR (treatment effect, *F*_1,451_ = 0.7, *P* = 0.4). Compared to WT, HR mice exhibited an overall ~3% higher startle response (genotype effect, *F*_1,451_ = 7.2, *P* = 0.008), which varied by the acoustic intensity of the pulse (**Fig. 5B**, genotype x db interaction, *F*_5,408_ = 2.8, *P* = 0.016). The increased startle tended to be stronger at 120db (**Fig. 5B**, paired *t* test, t = 2, *P_adj_* = 0.08 marginally significant).

**Figure 5:**
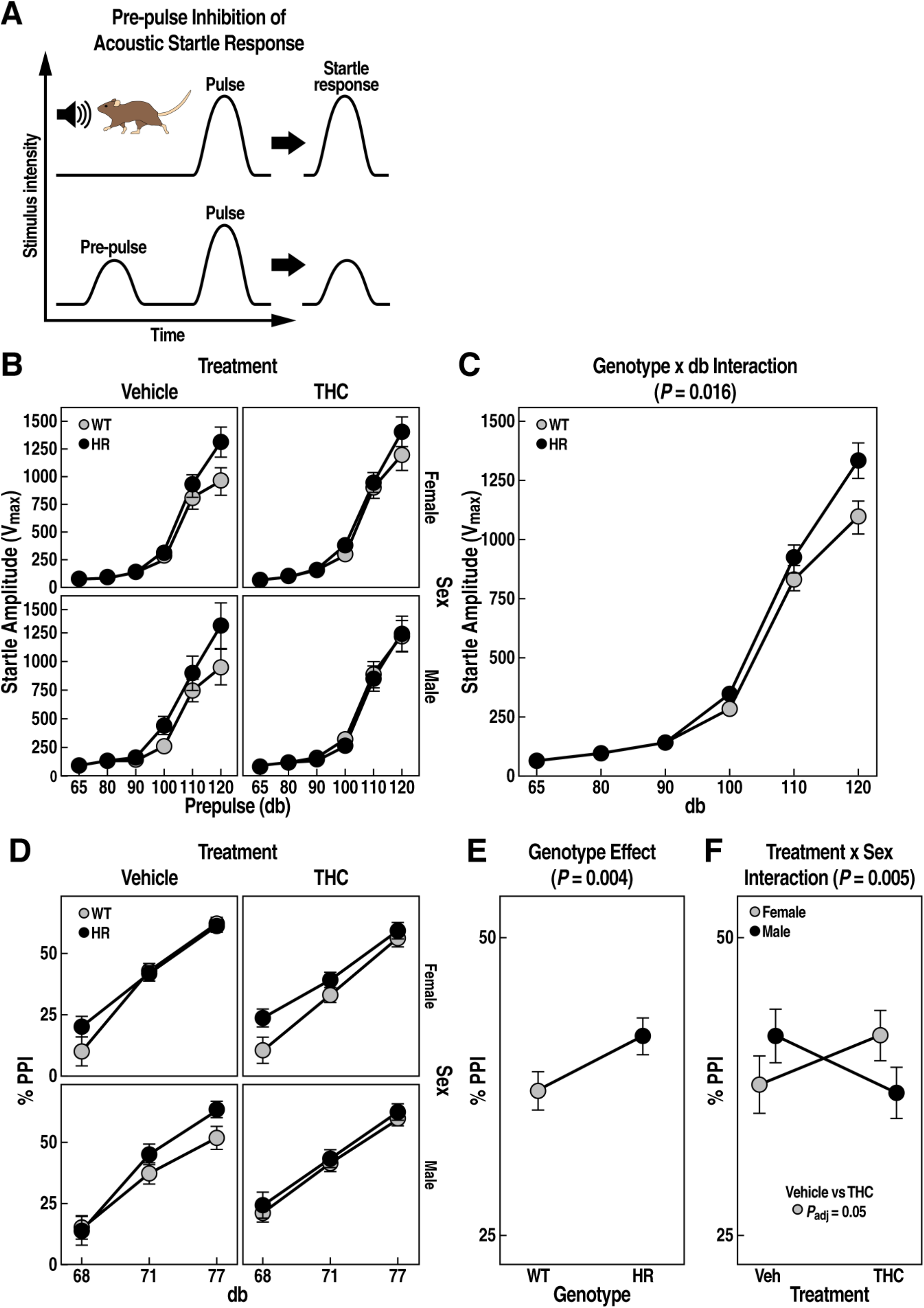
Acoustic startle reflex of pre-pulse inhibition in WT and HR mice following chronic adolescent exposure to THC. **(A)** Pre-pulse inhibition of acoustic startle test diagram. **(B)** Startle amplitude is expressed as Vmax ± SEM. **(C)** HR mice show increased startle response (genotype x db interaction, *P* = 0.016, LMM). **(D)** Pre-pulse inhibition is expressed as % PPI ± SEM. **(E)** HM mice exhibited higher % PPI compared to WT controls (main genotype effect, *P* = 0.004, LMM). **(F)** Female treated with THC during adolescence showed reduced %PPI compared to vehicle (treatment x sex interaction, *P* = 0.005, LMM). Adjusted *P* < 0.05 for planned post-hoc pairwise t tests are reported.

When examining the % PPI at three pre-pulse intensities (**Fig. 5C**, 68, 71 and 77 db), HR mice showed an overall ~13% increase in % PPI compared to WT mice (**Fig. 5D**, genotype effect, *F*_1,482_ = 8.6, *P* = 0.004). Adolescent exposure to THC also had an effect on % PPI that varied among sexes (**Fig. 5E**, treatment x sex interaction, *F*_1,482_ = 8, *P* = 0.005). Female mice treated with THC tended to show ~12 % reduction in % PPI compared to vehicle-treated mice (paired *t* test, t = 2, *P_adj_* = 0.05 marginally significant).

### THC and Reelin deficiency influenced behavioral domains that were associated with cognition and disinhibition

We performed a factor analysis to determine whether the multiple behavioral responses observed in our study reflect a smaller number of synthetic behavioral domains that are influenced by THC and *Reelin* deficiency.

The factor analysis identified three unique behavioral domains (factors), which explained 57% of total variance (**Fig. 6A**). The first factor explained 23.8% of the total variance and contained outcomes (positive loadings) from the social interaction and working memory tests (e.g. time sniffing novel mouse, discrimination index). As factor 1 largely reflected measures of cognitive functions, we named it “*cognition*”. The second factor (17% of the total variance) was named “*disinhibition*” as it contained behaviors that reflected increased exploratory or disinhibited behavior (e.g. time in the center of the open field, novel object exploration). The third factor explained 15.7% of the total variance and was named “*stress reactivity*” because the main outcome that contributed to this cluster was the mobility time of the TS test.

**Figure 6:**
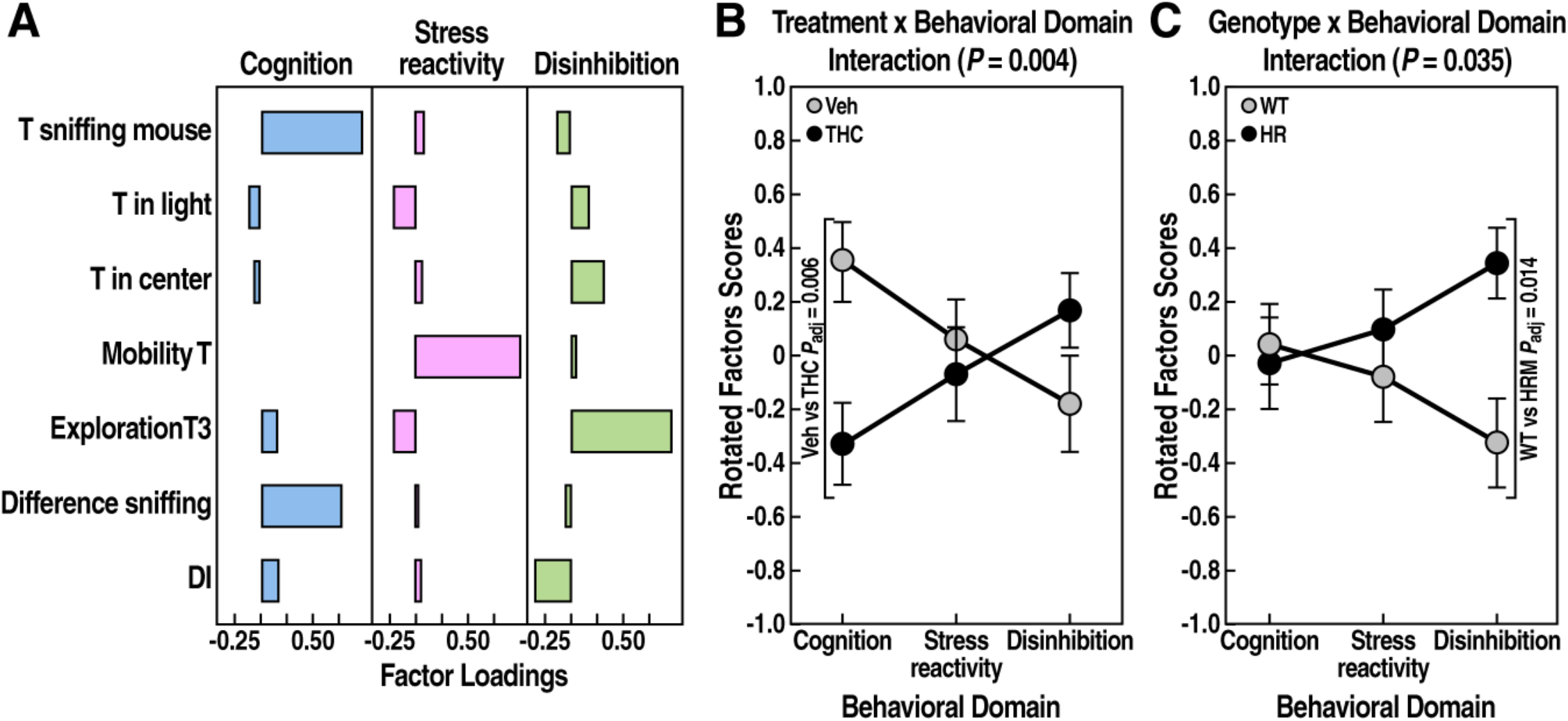
Factor analysis of behavioral tests. **(A)** Factor loadings for each named behavioral domain are reported for several behavioral variables. LMM analysis for rotated factor scores revealed a **(B)** treatment x behavioral domain interaction (*P* = 0.04), and a **(C)** genotype x behavioral domain interaction (*P* = 0.035). Adjusted *P* < 0.05 for planned post-hoc pairwise *t* tests are reported.

To determine whether these behavioral domains differed between treatments, genotypes, or sexes, we analyzed the rotator factor scores for each subject (**Fig. 6B-C**). We found that the behavioral domains were significantly influenced by treatment or genotype (treatment x behavioral domains interaction, *F*_2,142_ = 5.7, *P* = 0.004; genotype x behavioral domains interaction, *F*_2,142_ = 3.4, *P* = 0.035; respectively). Post-hoc analysis revealed that *cognition* was significantly decreased by THC (**Fig. 6B**, paired *t* test, t = 3.1, *P_adj_* = 0.006), and *disinhibition* was significantly increased by *Reelin* deficiency (**Fig. 6C**, paired *t* test, t = 2.8, *P_adj_* = 0.014). Neither treatment, genotype, nor sex significantly influenced the third behavioral domain, *stress reactivity*.

Overall, this exploratory data analysis is consistent with the results of the separate behavioral tests by indicating that adolescent exposure to THC detrimentally affected cognitive functions, and it revealed that reduced expression of *Reelin* led to disinhibitory behaviors, which were intensified by THC treatment in a sex-specific manner.

### Novelty seeking in HR mice was associated with reduced neuronal activity in the lateral septum

In response to different environmental stimuli, the expression of *Fos* is rapidly induced in specific neuronal populations allowing the identification of relevant neuronal ensembles encoding specific behaviors (Curran and Morgan, 1995; Chaudhuri, 1997). To explore the neuronal ensemble underlying the enhanced novelty seeking behavior observed in the HR mice (**Fig. 1E**), we mapped the regional brain expression of Fos protein by IHC in juvenile mice subjected to the 6-DOT (**Fig. 1A**).

In response to this behavioral task, WT mice activated the expression of Fos in four brain areas, including mPFC, LS, DG and CA1 sub-regions of the HP (**Suppl. Fig. 4**). Thus, we selected these areas for measuring the number of Fos-expressing (Fos+) cells in WT and HR mice, thirty minutes after the end of the task (**Fig. 7A-B**). There were no significant differences in Fos expression between genotypes or sex in the mPFC, DG or CA1 (**Fig. 7F,H-I**). In contrast, compared to WT, HR mice showed a ~28% reduction in number of Fos+ cells in the LS (**Fig. 7G**, genotype effect, *F*_1,13_ = 5, *P* = 0.04). To examine the expression of *Reelin* and *Fos* in the LS, we used RNA FISH and showed that *Reelin*-expressing and *Fos*-expressing cells are predominately located in the LS caudal (LSc), with fewer positive cells present in the LS rostral (LSr) (**Fig. 8A,C**). Consistent with the expression of *Reelin* by GABAergic neurons (Lossi et al., 2019), *Reelin*+ cells also co-expressed the vesicular GABA transporter (*Scl32a1*), a marker of GABAergic neurons (**Fig. 8B**). To further characterize the *Fos*-positive ensemble, we co-labeled *Fos*+ cells with markers of GABAergic (*Scl32a1*) and cholinergic neurons (*Chat*), which represent the major neuronal cell types of the LS. We found that *Fos* transcript was activated in both cell types and that the location of *Fos*+ cells showed a preferential location in the LSc, similarly to those expressing *Reelin* (**Fig. 8D**).

**Figure 7:**
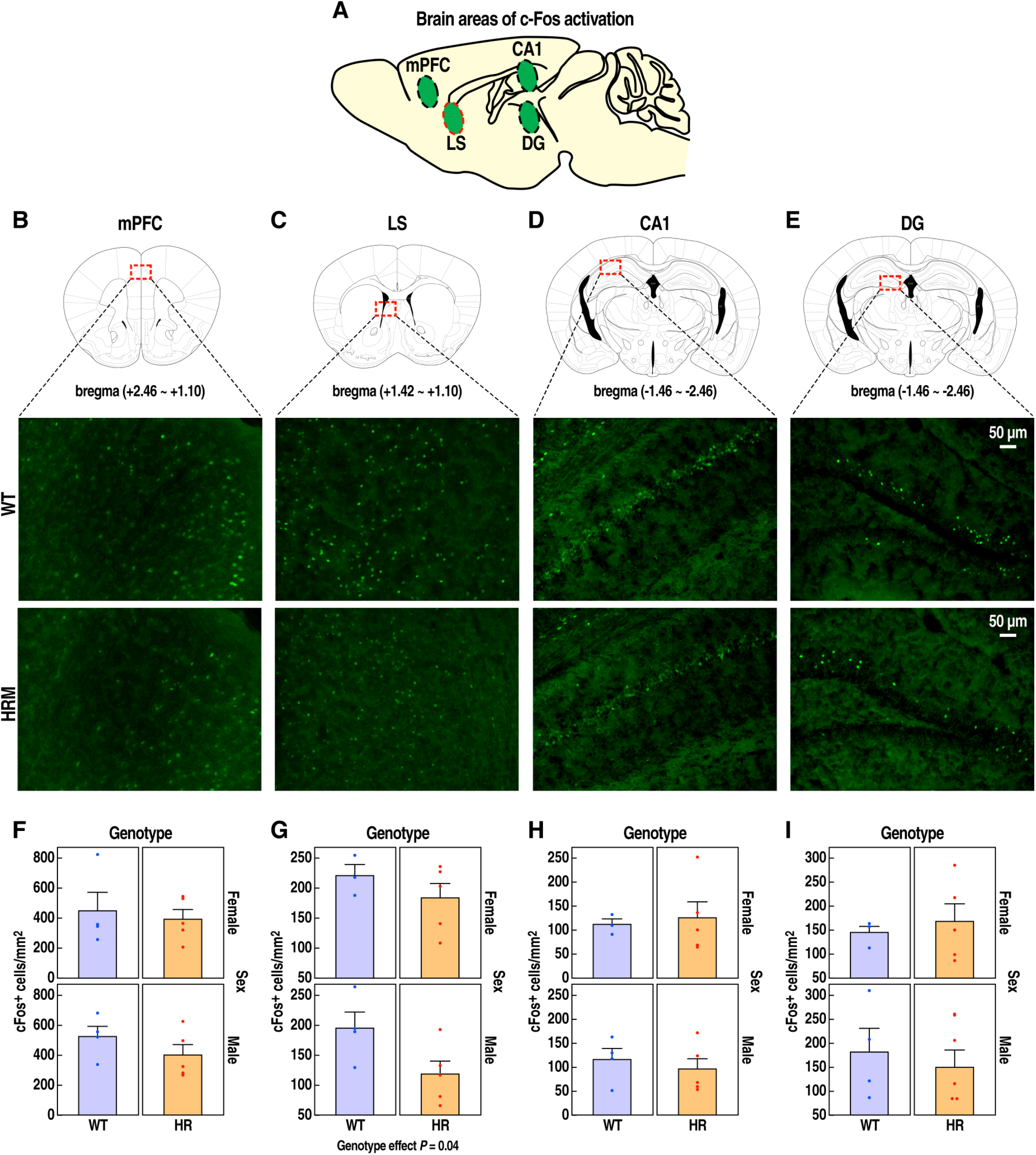
Fos expression is reduced in the LS of HR mice in response to novel object exploration. **(A)** Brain schematic illustrating the target areas for Fos quantification: mPFC, LS, CA1, DG. (**B-E**) Schematic brain palette with bregma levels (top) and representative images (bottom) of the mPFC, LS, CA1, and DG in WT (*n* = 6 male, *n* = 4 female) and HR mice (*n* = 6 male, *n* = 9 female). Scale bar, 50um.

**Figure 8:**
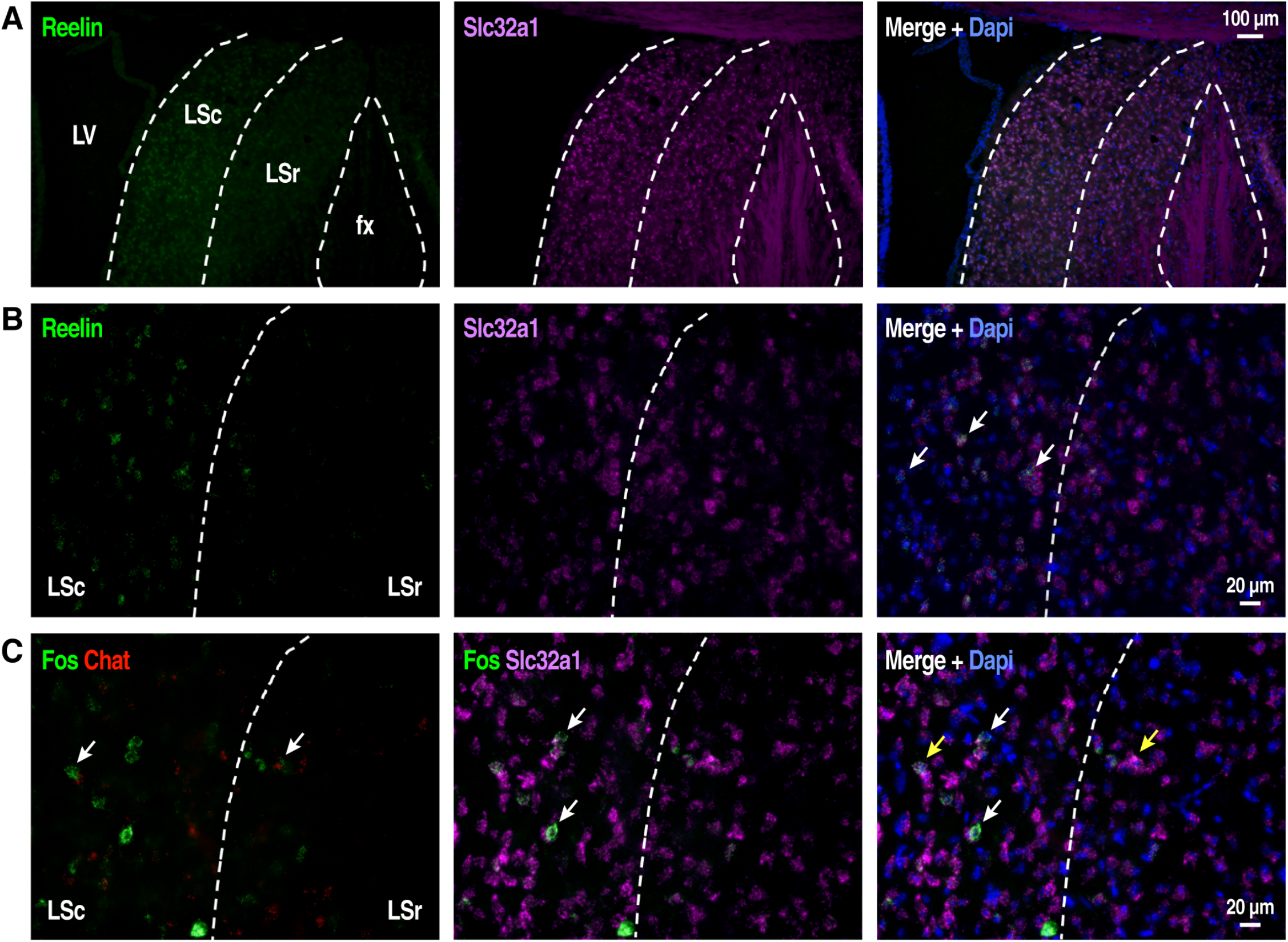
Distribution of different cell types of the the LS. **(A)** RNA FISH showing cells expressing *Reelin* and *Slc32a1* relative to Dapi-stained nuclei in the LS at bregma +0.50 mm. **(B)** Higher magnification images illustrate the co-localization (white arrow) of *Reelin* and *Slc32a1* mRNA. **(C)** RNA FISH showing cells expressing *Fos* and *Slc32a1* relative to Dapi-stained nuclei in the LS at bregma +0.26 mm. **(D)** Higher magnification images illustrate the co-localization of *Fos* and *Slc32a1* (white arrow) or *Fos* and *Chat*) mRNA. Dotted lines indicate the caudal (LSc) and rostral (LSr) regions of the LS, and the fornix (fx) fibers. Scale bars are 100μm in **(A)** and 20μm in **(B-C)**.

## Discussion

The present study revealed for the first time that reduced levels of *Reelin* influences behavioral abnormalities caused by heavy consumption of THC during adolescence, and that Reelin plays a role in the neurobiological mechanisms underlying novelty seeking by modulating neuronal ensembles in the LS.

### Influence of THC and Reelin deficiency on cognitive functions

We found that chronic adolescent exposure to THC impaired working memory, in line with previous reports in rodent models and clinical studies in humans (Renard et al., 2014; Rubino and Parolaro, 2016; Hurd et al., 2019). Additionally, HR mice showed lower working memory in comparison to the WT littermates and to a similar extent as did chronic adolescent exposure to THC. Although it is the first time that HR mice are tested in a task that measures memory span, such as the 6-DOT, this finding is consistent with studies demonstrating the crucial role of Reelin in mechanisms underlying learning and memory (Weeber et al., 2002; Rogers et al., 2013; Iafrati et al., 2014; Telese et al., 2015).

We found that social interaction was impaired by chronic adolescent exposure to THC with the strongest effects in HR mice. These observations are in line with evidence in human and preclinical studies showing social deficits induced by THC (Long et al., 2010). Only few studies have examined social behaviors of the HR mice and did not report prominent social deficits (Podhorna and Didriksen, 2004; Macri et al., 2010; Michetti et al., 2014). Thus, our study revealed for the first time that *Reelin* deficiency exacerbates social deficits induced by THC.

Consistently, the factor analysis revealed that variables from both working memory and social interaction tests contributed to a behavioral domain, named ‘*cognition*’, which was negatively influenced by adolescent exposure to THC.

### Reelin deficiency is associated with disinhibitory phenotypes

We speculate that multiple behavioral responses exhibited by HR mice may reflect general behavioral disinhibition. In rodents, behavioral disinhibition is measured as a function of increased exploratory activity towards unfamiliar objects or environments, and has been associated with compulsive drug taking (Davis et al., 2008; Flagel et al., 2010; Belin et al., 2011). In humans, behavioral disinhibition reflects personality traits that encompasses impulsivity, risk-taking and novelty seeking phenotypes; these behavioral patterns are more pronounced in adolescence and have been linked to addiction susceptibility (Young et al., 2009). Here, we found that, compared to WT, both male and female HR mice explored the novel objects for longer time in the 6-DOT. Additionally, female HR mice treated with THC showed signs of reduced inhibitory control as they spent more time in the center chamber of the OF test or in the brightly lit compartment of the LD test. Consistently, variables from these tests contributed to the same behavioral domain (‘*disinhibition*’) revealed by the factor analysis, which was positively influenced by *Reelin* deficiency. These observations are also in line with a previous study showing decreased anxiety of HR mice in the elevated plus maze and increased motor impulsivity (Ognibene et al., 2007).

Collectively, these results raise the hypothesis that impairment in Reelin signaling may underlie addiction traits by disrupting inhibitory control. Further supporting this hypothesis, in a recent genome-wide association study (GWAS), a variant of the *Reelin* gene was associated with higher likelihood to consume alcohol (*P* = 4 x 10^−9^, variant and risk allele rs756747-T) (Karlsson Linner et al., 2019).

### Reelin deficiency leads to abnormal responses to aversive conditions

We observed abnormal behavior in HR mice in response to aversive conditions. First, male HR mice treated with THC showed prolonged mobility time in the TS test compared to vehicle group. This behavior has been linked to proactive coping in response to a threat (Strekalova et al., 2004; Brockhurst et al., 2015). Second, HR mice exhibited enhanced startle reactivity to acoustic stimuli compared to WT mice, which was not influenced by THC treatment. In a previous study, the startling response of HR mice did not change following a single acoustic pulse of 105db; however, in our study, we used five startling pulses from 80db to 120db that likely increased the sensitivity of the task. Elevated ASR has been observed in mental conditions associated with impaired emotional reactivity, such as posttraumatic stress disorders (Orr et al., 1995) and obsessive-compulsive disorders (Kumari et al., 2001). These observations led us speculate that reduced expression of *Reelin* may lead to altered emotional reactivity. This hypothesis deserves further examination of the HR mice using tasks specifically designed to assess impulsivity and aggression.

### Reelin as a susceptibility factors for psychiatric disorders

The array of behavioral phenotypes exhibited by HR mice in our study, encompassing memory impairments, social deficits, poor inhibitory control, and altered stress responses, is reminiscent of the behaviors characterizing numerous psychiatric conditions, ranging from autism spectrum disorders (ASD) to schizophrenia and substance use disorders. Consistently, a role of Reelin in the development of these disorders is supported by several lines of evidence. In particular, whole exome sequencing studies identified *de novo* mutations in the *Reelin* gene in individuals with ASD (Neale et al., 2012; Iossifov et al., 2014; Wang et al., 2014). Further support for Reelin involvement in psychiatric disorders is provided by the observation of reduced expression of *Reelin* transcript or protein in postmortem brains of individuals affected by ASD (Fatemi et al., 2001), and schizophrenia (Guidotti et al., 2000; Ruzicka et al., 2007; Habl et al., 2012). These observations suggest that altered Reelin signaling may be a vulnerability factor for psychiatric disorders and that the HR mice represent a valuable animal model for translational research. However, we did not observe any PPI deficits in HR mice. In contrast, a previous study showed that a single *in vivo* injection of Reelin protein increases % PPI (Rogers et al., 2013). It is possible that, in our study, the elevated startle response observed in HR mice may act as a confounding factor for accurately assessing the effect of *Reelin* deficiency. It is also possible that impaired Reelin signaling contributes mainly to the cognitive, social and emotional deficits associated with schizophrenia, which are referred to as negative symptoms, as opposed to positive symptoms that include psychotic-like behaviors (Kirkpatrick et al., 2006).

Notably, THC reduced % PPI in female WT mice, suggesting that a history of drug exposure leads to psychotic-like behaviors. Conflicting results have been reported concerning the effects of adolescent chronic exposure to THC on psychotic-like behaviors in rodents (Rodriguez et al., 2017; Todd et al., 2017; Ibarra-Lecue et al., 2018). Differences in THC doses, mice genetic background and specific experimental conditions among different research groups may explain some of these conflicts.

### Sex differences in behavioral responses associated with Reelin haploinsufficiency

Our study revealed numerous sex differences in the behavioral abnormalities associated with *Reelin* deficiency and/or adolescent exposure to THC. Male HR mice were more sensitive to working memory impairments, as well as to emotional reactivity in the TS test. In contrast, female mice, showed reduced anxiety-like behaviors (HR) or reduced % PPI (WT) in response to THC. Consistently, previous finding in rodents showed that chronic adolescent exposure to cannabinoids is associated with numerous behavioral sex differences; however, the knowledge of the underlying mechanisms remains limited (Craft et al., 2013). Given the critical role of Reelin brain development, it is possible that Reelin influences the development of the endocannabinoid system in a sex-specific manner, which can lead to sex differences to the behavioral effects of THC.

### The novelty seeking phenotype observed in HR mice is associated with reduced Fos expression in the LS

We found that the prolonged exploratory activity of novel objects exhibited by HR mice in the 6-DOT was associated with reduced neuronal activity in the LS. Although the LS has not been linked directly to novelty seeking, it has been implicated in the modulation of rewarding experiences that are correlated to novelty seeking (Luo et al., 2011; Harasta et al., 2015; McGlinchey and Aston-Jones, 2018). Moreover, several behaviors influenced by *Reelin* deficiency in our study have also been linked to the activity of the LS, such as social interaction, memory and emotional processes associated with aversive situations (Singewald et al., 2011; Leroy et al., 2018) (Shin et al., 2018). Notably, *Fos* and *Reelin* expression are detected in the LSc, a region implicated in the regulation of drug-seeking behavior via a population of GABAergic neurons that project to the ventral tegmental area (VTA) (Luo et al., 2011). We speculate that *Reelin* deficiency may alter LS-VTA pathway underlying motivated behavior by disrupting GABA inhibition in the LSc and, hence, disinhibiting dopaminergic neurons in the VTA. In our previous work, we showed that Reelin triggers a signaling cascade via the intracellular domain of its receptor LRP8 and directly activates transcriptional enhancers of immediate early genes, including *Fos* (Telese et al., 2015). Thus, it is possible that Reelin signaling modulates the transcriptional and epigenomic landscape of specific neuronal populations in the LS.

In conclusion, our study suggests that elucidating Reelin signaling will improve our understanding of neurobiological mechanisms underlying behavioral traits relevant to the development of psychiatric conditions.

## Acknowledgments

This work was supported by the National Institute on Drug Abuse [DP1DA042232 to FT]. We thank R.F. Hechinger and S. Roige-Sanchez for helpful discussions and critical reading of the manuscript; A. Turner, H. Taylor, S. Nolan, A. Apelian, and J. Hightower for technical assistance.

---

**Supplementary Figure 1:**
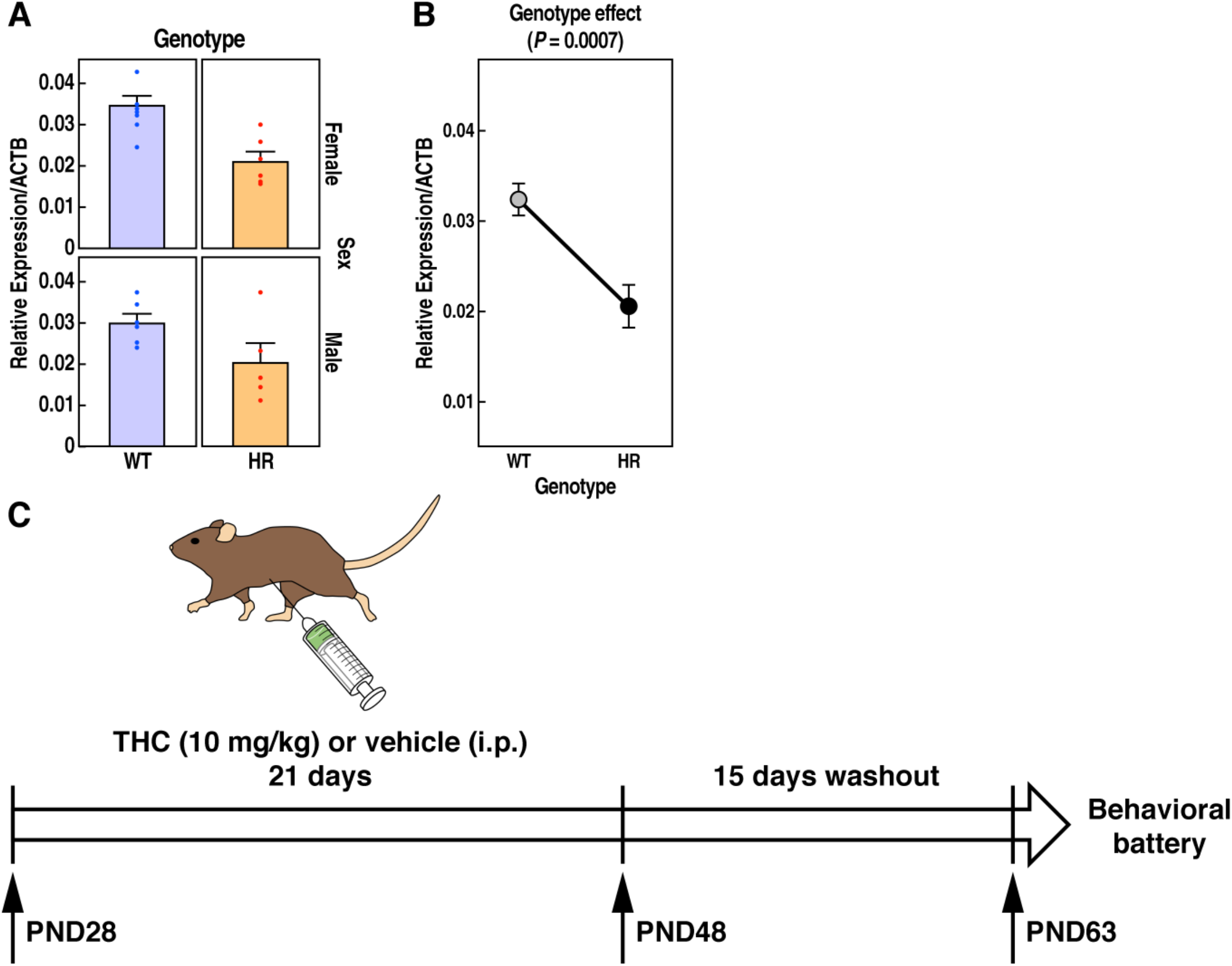
HR mice are used for chronic adolescent administration of THC. **(A)** RT-PCR indicate Reelin mRNA expression in brain tissues isolated from WT and HR mice. Level of Reelin expression are normalized with the housekeeping gene ACTB. **(B)** Reelin mRNA is reduced in HR mice compared to WT littermates (genotype effect, *P* = 0.0007, LMM). *n* = 8 female WT, *n* = 6 male WT, n = 6 female HR, *n* = 5 male HR mice. **(C)** Schematic diagram illustrates the daily i.p. administration of THC (10 mg/Kg) during adolescence (PND28-48), and the day when behavioral assessment began (PDN63) after 15 days of washout.

**Supplementary Figure 2:**
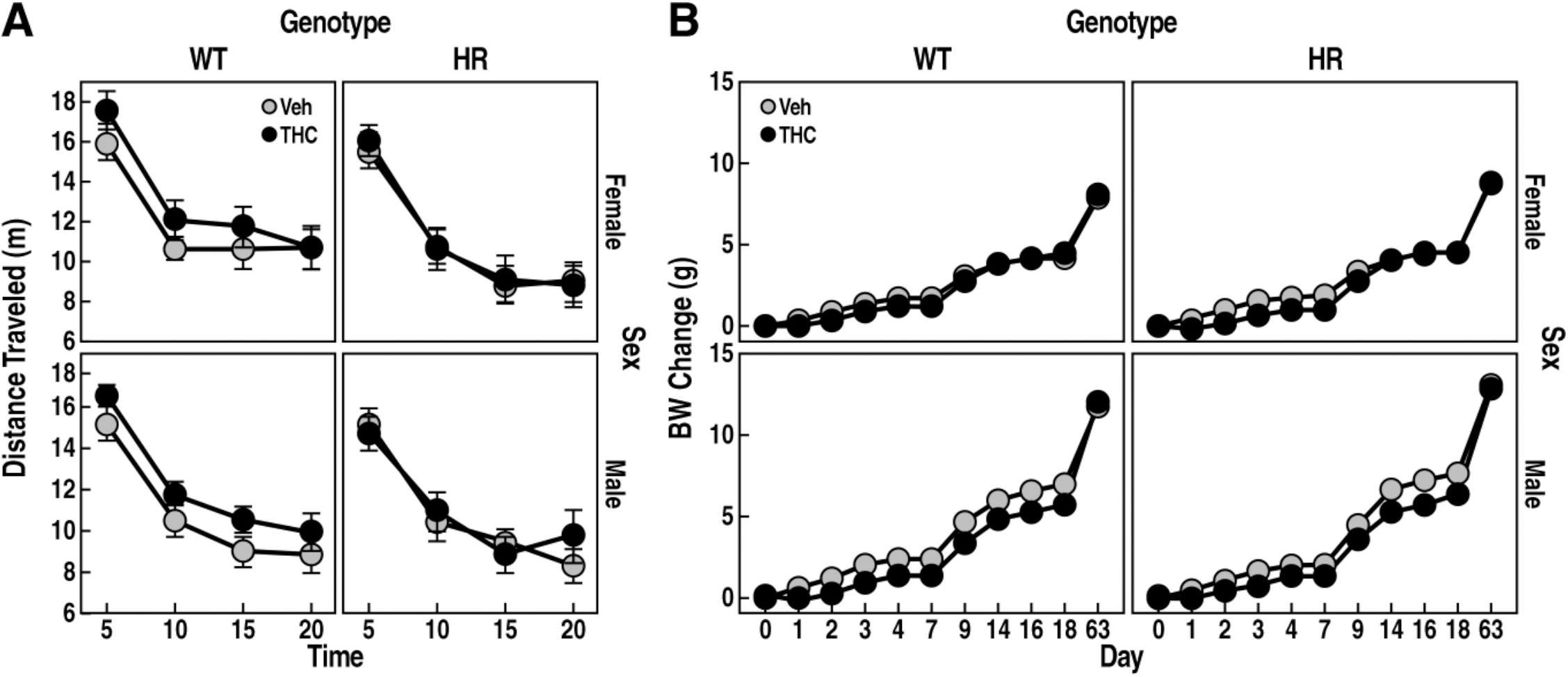
Locomotor activity and body weight for WT and HR mice. **(A)** Twenty-minute locomotor activity is expressed as mean distance traveled (m) ± SEM for every 5-minute intervals. **(B)** Body weight (g) change is expressed as mean ± SEM for the THC administration course (day 1 to 14), and for day 63 when the behavioral testing started.

**Supplementary Figure 3:**
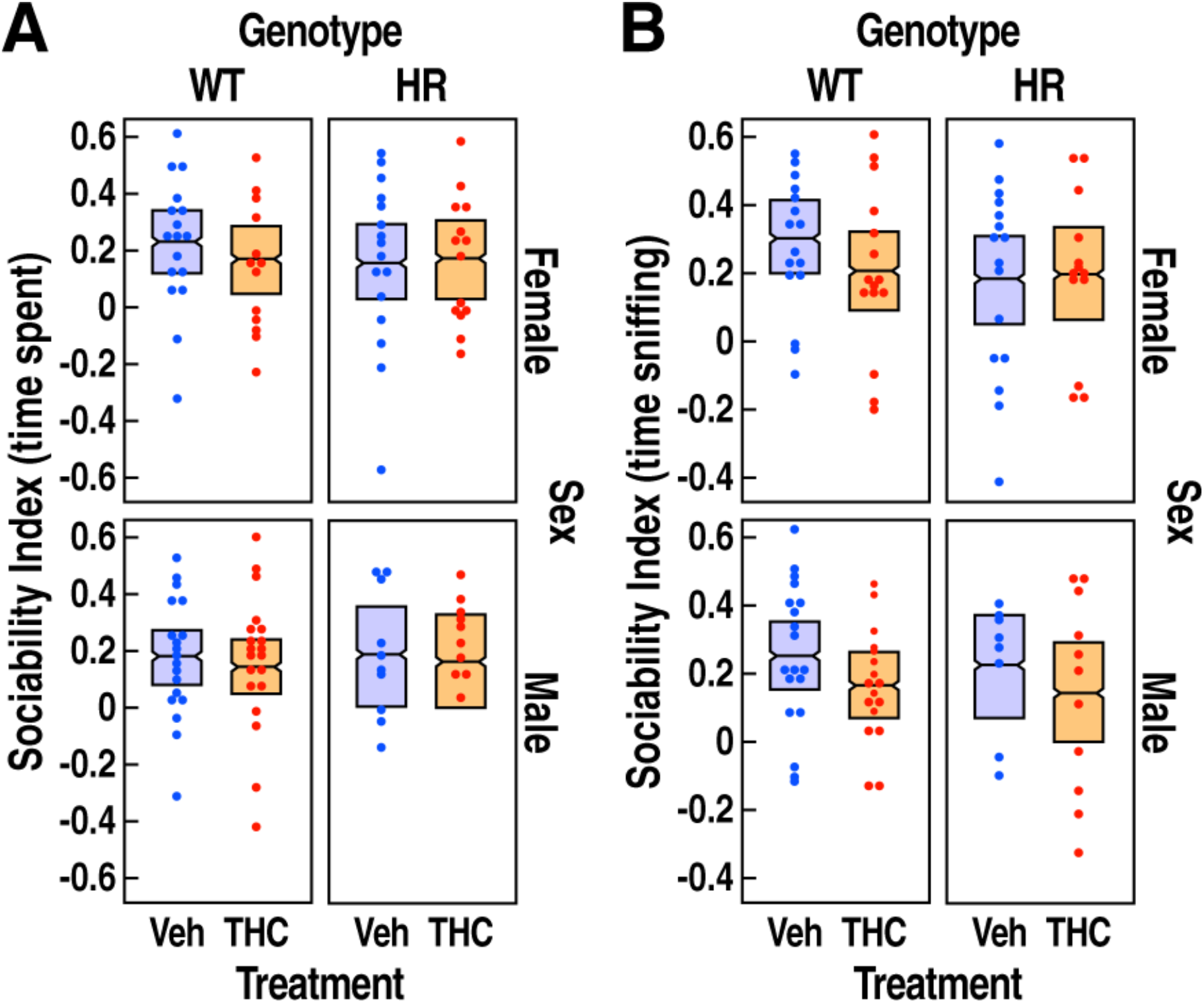
Sociability index in WT and HR mice following chronic adolescent exposure to THC. **(A)** Sociability index calculated using time spent in the chamber with novel mouse versus novel object is shown as mean ± 95% confidence intervals. **(B)** Sociability index calculated using time sniffing novel mouse versus novel object is shown as mean ± 95% confidence intervals. Treatment, genotype and sex had no significant effect on both sociability indices.

**Supplementary Figure 4:**
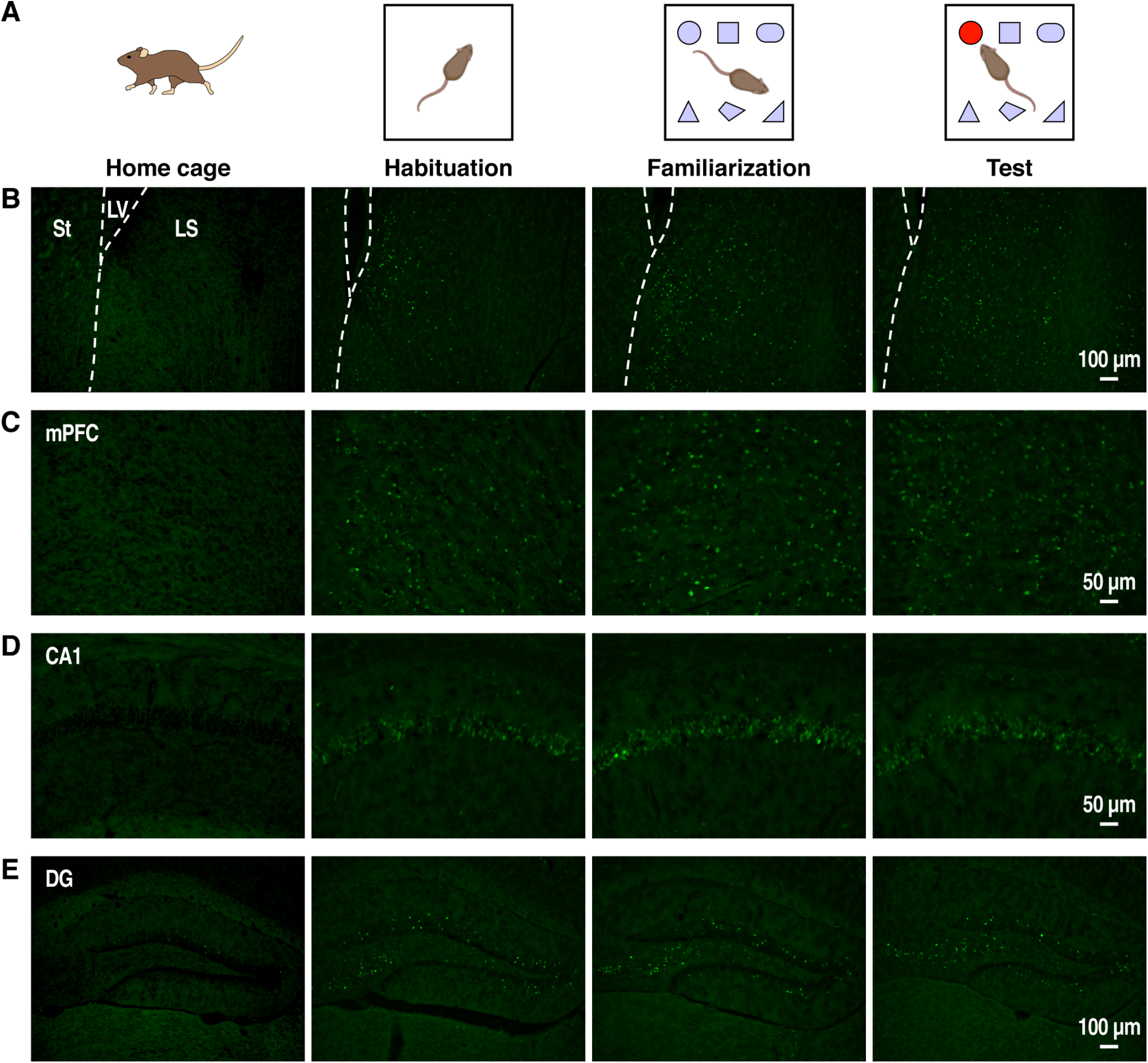
Fos activation in response to novelty. **(A)** Schematic of behavioral manipulation, including home cage (control), habituation, familiarization, and test phases of the 6-DOT. **(B-C)** Representative images of Fos expression by IHC in different brain regions, including LS, mPFC, CA1 and DG. Dotted lines illustrate the line separating the LS from the striatum (St) and the lateral ventricle (LV). Scale bars are 100μm in **(B, E)** and 50μm in **(C, D)**.

